# Competitive SNP-LAMP probes for rapid and robust single-nucleotide polymorphism detection

**DOI:** 10.1101/2021.03.29.437576

**Authors:** Leland B. Hyman, Clare R. Christopher, Philip A. Romero

**Affiliations:** Department of Biochemistry, University of Wisconsin--Madison, Madison, WI, USA; Department of Chemical & Biological Engineering, University of Wisconsin--Madison, Madison, WI, USA; The University of Wisconsin Carbone Cancer Center, Madison, WI, USA

## Abstract

Single-nucleotide polymorphisms (SNPs) are the most common source of genetic variation between individuals and have implications in human disease, pathogen drug resistance, and agriculture. SNPs are typically detected using DNA sequencing, which requires advanced sample preparation and instrumentation, and thus cannot be deployed for on-site testing or in low-resource settings. In this work we have developed a simple and robust assay to rapidly detect SNPs in nucleic acid samples. Our approach combines LAMP-based target amplification with fluorescent probes to detect SNPs with high specificity in a one-pot reaction format. A competitive “sink” strand preferentially binds to off-target products and shifts the free energy landscape to favor specific activation by SNP products. We demonstrated the broad utility and reliability of our SNP-LAMP method by detecting three distinct SNPs across the human genome. We also designed an assay to rapidly detect highly transmissible SARS-CoV-2 variants. This work demonstrates that competitive SNP-LAMP is a powerful and universal method that could be applied in point-of-care settings to detect any target SNP with high specificity and sensitivity.

## Introduction

Single-nucleotide polymorphisms (SNPs) are the most widespread source of genetic variation between individuals^1^. A single base substitution can induce profound changes in a protein’s structure, altering its enzymatic function^2^, cellular trafficking^3^, or solubility^4^. As a result, SNPs play crucial roles in many different biological phenomena including human^5^ and animal disease^6^, pathogen drug resistance^7^, and agricultural blight^8^. SNPs are routinely detected using DNA sequencing^9^, which requires a laboratory setting for sample preparation, in addition to large, expensive, and slow DNA sequencing instruments. Current DNA sequencing methods cannot be adapted to low-resource settings such as rural areas or developing countries^10^. There is a substantial need for rapid and point-of-care SNP detection assays that can be performed on-site without advanced laboratory equipment or a cold supply chain.

Loop-mediated isothermal amplification (LAMP) is a simple and robust method for sequence-specific detection of nucleic acids^11–13^. Unlike PCR, the amplification process occurs continuously at a temperature of 65°C, facilitating fast amplification times and use of a simple heated block rather than a thermocycler^14^. Previous work has developed LAMP-based assays to detect SNPs. Mismatch SNP-LAMP incorporates the SNP base into the 3’ terminus of a LAMP primer, causing a mismatch and preventing polymerase extension when the non-SNP sequence is present^15–17^. However, heterogeneity in primer synthesis and promiscuous mismatch extension by the LAMP polymerase^18,19^ can lead to unpredictable amplification times and false positive events. Other SNP-LAMP strategies use fluorescent DNA probes to detect SNPs in LAMP products^20^. The probe consists of a DNA duplex with a quenched fluorophore that is complimentary to the SNP sequence and thus preferentially binds the SNP over the non-SNP. The difference in binding energies from a single mismatched base can be quite small, leading to substantial signal from non-SNP sequences that is difficult to distinguish SNP sequences^21^.

We have developed a novel SNP-LAMP method that can rapidly and robustly detect SNPs in a simple, one-pot assay format. Our approach leverages LAMP-based target amplification and competitive fluorescent probes^21–24^ to specifically detect SNP over non-SNP sequences. Competitive “sink” strands preferentially bind off-target sequences and help to widen the free energy gap between highly similar SNP and non-SNP sequences. We devised a thermodynamics-based computational optimization algorithm to design probe sets with high sensitivity and specificity for a target SNP. We demonstrated the ability of competitive SNP-LAMP to detect specific SNPs from highly complex total RNA samples in a simple one-pot reaction. Finally, we developed a simple and streamlined assay to detect SARS-CoV-2 strains that could be used to monitor emerging variant outbreaks. Competitive SNP-LAMP is a powerful and robust solution for detecting SNPs that is simple and inexpensive enough to be deployed on a large scale and in low-resource settings.

## Results

### Competitive probes for highly specific LAMP-based detection of SNPs

We sought to identify a robust isothermal approach to detect SNPs in diverse nucleic acid samples. Previous work has found the signal of strand displacement probes can be enhanced by including a ‘sink’ complex that competes for binding with the non-SNP sequence^22–24^. This competitive probe alters the free energy landscape and has achieved remarkable specificity for PCR-based SNP detection^21^. We adapted this approach to detect SNPs in LAMP products.

Our system includes a strand displacement probe duplex that is complementary to the SNP sequence and a competitive sink duplex that is complementary to the non-SNP sequence (Fig. 1ab). Non-SNP LAMP products will preferentially bind to the sink strand over the fluorescent probe, and reduce the off-target signal. As a proof of concept, we first designed a set of probe and sink duplexes to detect the c.776A>C mutation in the human *ACTB* gene. Our competitive probe system could clearly distinguish non-SNP versus SNP samples (*p* = 1.6 × 10^−7^) (Fig. 1c). The fluorescence signal of the SNP sample was 2.7 times higher than the non-SNP sample. Additionally, the non-SNP signal was only 15% higher than the non-template control, indicating that the sink complex was successfully reducing off-target binding. We also performed the reaction without the sink complex and found that both the SNP and non-SNP targets activated, with no significant difference in signal between the two (p=0.46). These results suggested that competitive fluorescent probes are suitable for SNP detection within single-stranded LAMP amplicon regions.

**Figure 1.**
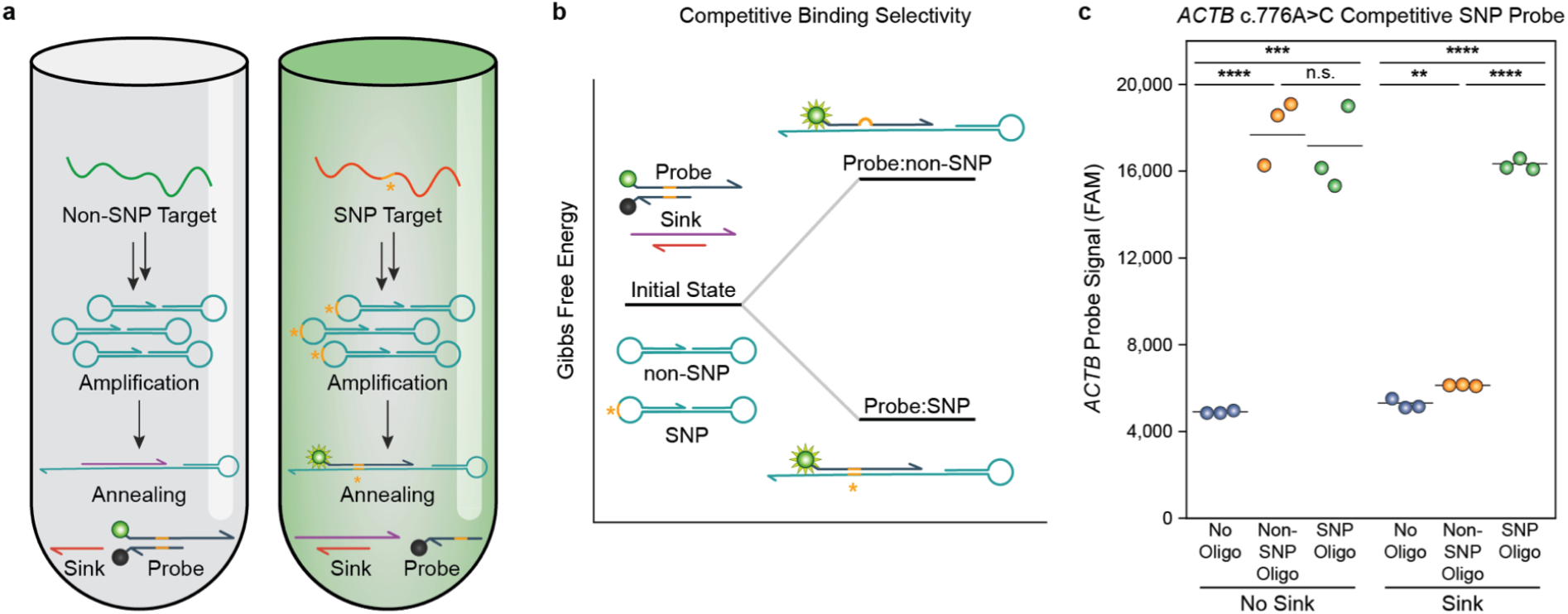
A competitive SNP-LAMP detection strategy. (**a**) Schematic of the competitive SNP-LAMP assay. SNP and non-SNP targets are amplified in a standard LAMP reaction, after which the amplicons are melted and annealed to a SNP-specific probe and a non-SNP-specific sink complex. The probe strand predominantly binds to the SNP sequence, producing a fluorescent signal. (**b**) SNP-LAMP competitive probe strategy and thermodynamics. In addition to the probe complex, a non-SNP-specific ‘sink’ complex is added, which competes for non-SNP binding and greatly increases specificity. This is reflected in a large ΔG difference between the Probe:SNP and Probe:non-SNP duplexes at equilibrium. (**c**) Detection of a c.776A>C mutation in the *ACTB* gene using competitive SNP-LAMP. Adding a sink complex greatly reduced the signal from a representative non-SNP oligo target (p=8.3*10^−5^), while producing no significant change in signal from a SNP oligo target (p=0.65).

### Computational design of competitive SNP-LAMP probes

Our competitive probe system consists of four DNA strands that can be of different lengths and complementary to different regions surrounding the target SNP. Our initial probe design involved manually tweaking the oligonucleotide sequences to achieve the desired behavior. However, the full design space is massive and shifting a strand by even a single base can drastically alter a design’s specificity. These factors make it challenging to manually design competitive SNP probes with optimal signal and specificity.

We developed a computational framework to design competitive probe combinations that maximally discriminate between SNP and non-SNP targets. Probe design is a multi-objective optimization problem that must balance two potentially conflicting goals: high fold activation in the presence of the target SNP and high SNP specificity over the non-SNP. We use thermodynamic modeling^25^ to evaluate how a given design will behave in the absence of any target, in the presence of the SNP target, and in the presence of the non-SNP target. These simulations provide an estimate of the fluorescence signal (concentration of unquenched probe) produced under these three conditions. We define an aggregate objective function that captures the two design objectives:

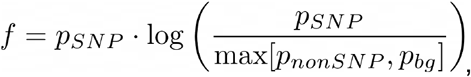

where *p*_*SNP*_, *p*_*nonSNP*_, and *p*_*bg*_ are the proportion of unquenched probe in SNP, non-SNP, and background (no template) thermodynamic simulations, respectively. The *p*_*SNP*_ term represents the fluorescence signal produced by the SNP, while the second term captures the difference between target and off-target signals. We seek to maximize this aggregate objective over all possible probe designs.

Our system consists of a probe duplex that contains a fluorophore-quencher pair and a sink duplex. These four DNA strands are each specified by their number of complementary bases before and after the SNP base position (Figure 2a). In order to better understand the probe design space, we simulated 10,000 random probe sets, each specific to a randomly generated SNP target sequence between 25 and 40 bases long. We found optimized probe sets are incredibly rare, with only 0.45% of these random designs displaying a signal greater than 70% and specificity over 100 fold (Fig. 2b). We generated a set of 12 biophysical features describing each probe (Table S1) and performed principal component analysis (PCA) to visualize the design space (Fig. S1). The optimized probe designs fall within a specific region that occupies roughly one quarter of the total design space area.

**Figure 2.**
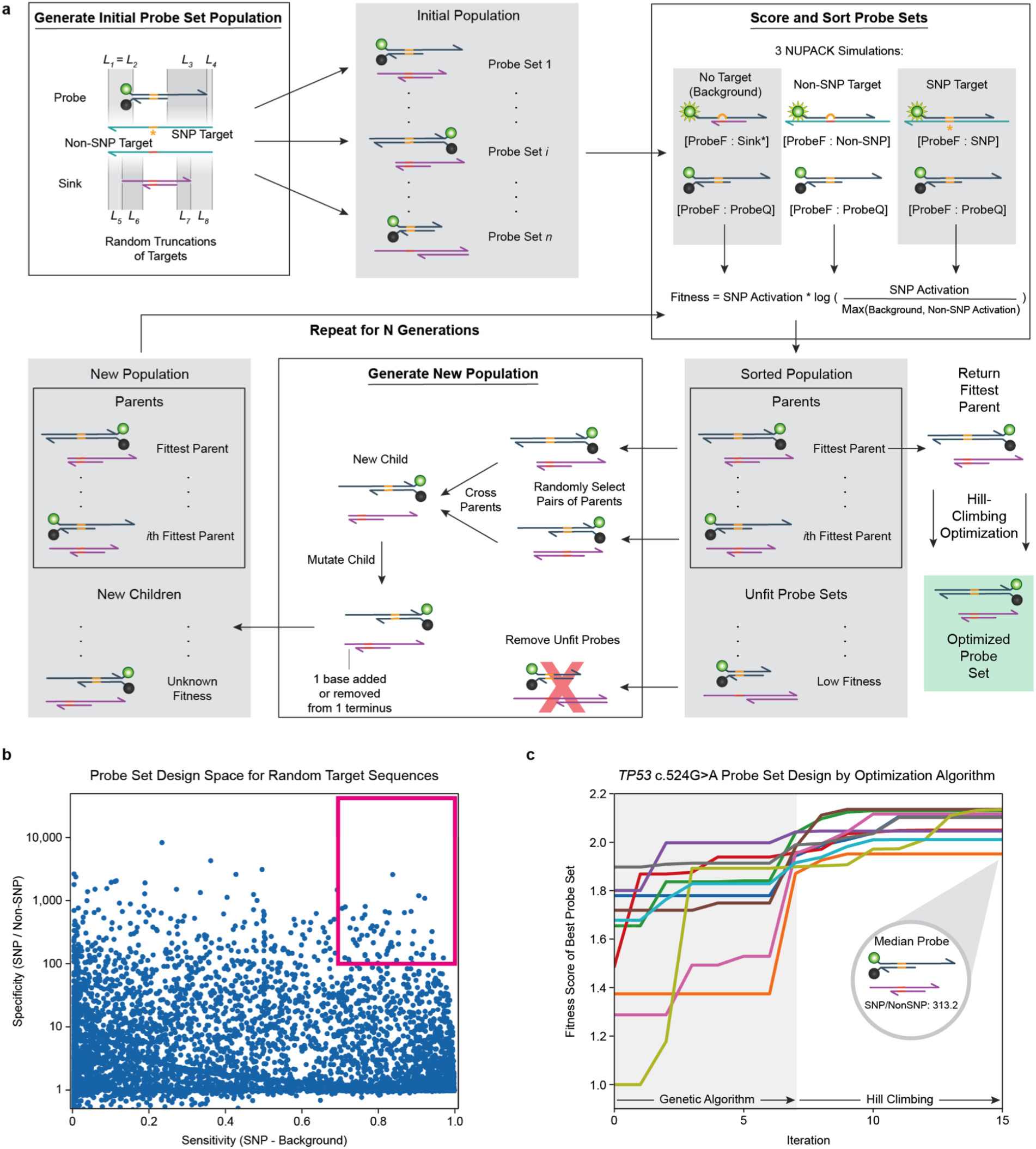
A hybrid genetic algorithm (GA) and hill-climbing optimization strategy for SNP-LAMP probe design. (**a**) During GA optimization, probes are randomly generated, mixed, and selected by fitness in a series of generations, mimicking natural evolution. After the designated number of generations, the fittest probe is further optimized by a hill-climbing algorithm to reach the nearest fitness maximum. (**b**) Design space for 10,000 random probes, each designed for a unique SNP target. High-fitness probes are rare: only 0.45% of designs fall within the optimal design space (magenta). (**c**) Convergence of the algorithm over 7 GA generations followed by hill-climbing. Across all 10 runs, the median optimal probes’ background-subtracted SNP/non-SNP specificity was consistently high, with a median of 313.2 and a standard deviation of 47.6.

We use a hybrid genetic algorithm-hill climbing algorithm to optimize our objective function over probe design space (Fig. 2a). The genetic algorithm searches the space broadly by iteratively breeding, mutating, and selecting top design candidates. The top sequence from the genetic algorithm is then optimized via hill climbing to exhaustively search the local design space and ensure we have reached a local maximum. The resulting probe design should balance signal and specificity for highly optimized for SNP detection. We tested our algorithm by designing a series of probe sets toward a c.524G>A mutation in the *TP53* tumor suppressor gene. We ran the algorithm 10 times from random starting points and observed its convergence to an optimal probe sequence (Fig. 2c). With only 7 GA generations and an initial population size of 128, the algorithm reliably converged to high specificity and high signal probe sets. Across all 10 runs, the median final probe design had a SNP/non-SNP specificity of 313.2 with a standard deviation of 47.6, and produced 85.5% of its maximum possible fluorescent signal when detecting the SNP sequence. Furthermore, the algorithm showed steady fitness improvement in nearly every case, suggesting that it can generate enough diversity to avoid becoming trapped in local minima. This algorithm should therefore serve as a reliable and effective means for competitive composition-based SNP probe design.

### One-pot competitive SNP-LAMP robustly detects SNPs in total RNA samples

We tested the generality and reliability of our computational design method by designing SNP-LAMP probes to detect three distinct SNPs across the human genome. The three targets include the c.186A>G mutation in the *MT-CO2* mitochondrial housekeeping gene, the c.524G>A mutation in the *TP53* tumor suppressor gene, and the c.4799T>C mutation in the *NOTCH1* oncogene. The SK-BR-3 and MOLT-4 human cell lines differ at these three targeted sites, and we used total RNA samples from each to test the performance of our designs.

We developed a one-pot SNP-LAMP assay where a nucleic acid sample is added to the five LAMP primers, the designed probe and sink duplexes, and a standard LAMP master mix (Fig. 3a). The sample is first incubated at 65 C for 60-75 minutes to allow LAMP-based target amplification, it is then heated to melt the LAMP products and slowly annealed to allow probe/sink hybridization to reach equilibrium. We performed our one-pot SNP-LAMP assay on each of the three targeted mutations and found it could reliably distinguish SNP template versus no template and SNP versus non-SNP template in all cases (Fig. 3b-d). We additionally verified through Sanger sequencing that each cell line’s RNA produced LAMP amplicons containing the expected SNP/non-SNP sequence (Fig. S2).

**Figure 3.**
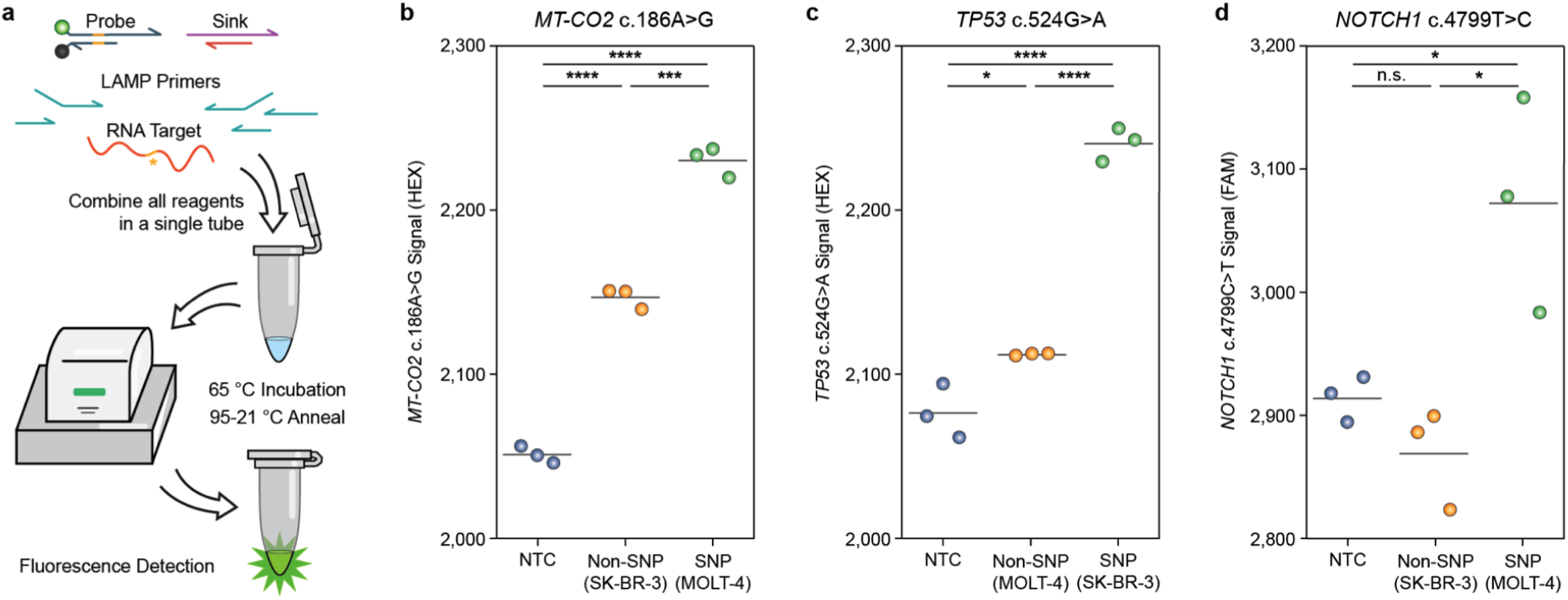
One-pot assay workflow and performance with total RNA as input. Endpoint fluorescence values are shown after LAMP and annealing. (**a**) Schematic of the one-pot SNP-LAMP assay workflow used in this work. LAMP primers, assay reagents, probe and sink strands, and RNA sample are added to a PCR tube, incubated and annealed on a standard qPCR machine, and measured for fluorescent signal. Once the tube is closed, no reagents are added or removed. (**b**) A competitive SNP-LAMP probe recognizing a c.186A>G mutation in the *MT-CO2* gene activated in response to MOLT-4 total RNA to produce a significantly larger signal than SK-BR-3 RNA (p=1.05*10^−4^) and background (p=3.96*10^−6^). (**c**) A competitive SNP-LAMP probe targeting a c.524G>A mutation in the *TP53* tumor suppressor gene. As expected, the *TP53* probe activated in response to SK-BR-3 total RNA, producing a significantly larger signal than MOLT-4 RNA (p=1.5*10^−5^) and background (p=6.4*10^−5^). (**d**) A competitive SNP-LAMP probe recognizing a c.4799T>C mutation in the *NOTCH1* oncogene activated in response to MOLT-4 total RNA to produce a significantly larger signal than SK-BR-3 RNA (p=0.011) and background (p=0.019).

To assess how our designed probe sets may perform with rare variant allelic frequencies, we performed experiments with varying proportions of SNP and non-SNP DNA oligos (Figs. S3a-b). The *TP53* and *NOTCH1* probes displayed a high sensitivity for the SNP at frequencies as low as 1%, with p-values of 2.3*10^−5^ and 3.1*10^−3^, respectively. This demonstrates that our competitive SNP probes can detect low proportions of SNP target in an overwhelming background of non-SNP sequences.

### A rapid test to distinguish SARS-CoV-2 variants

The COVID-19 pandemic has illustrated the importance of rapid point-of-care testing in disease mitigation and tracking^26^. All existing methods to monitor different SARS-CoV-2 strains involve DNA sequencing and cannot be easily deployed on a large scale or in low-resource settings^27^. This unmet need for low cost, rapid, and point-of-care strain tracking inspired us to develop a competitive SNP-LAMP test for specific SARS-CoV-2 variants.

We designed a probe set to target the D614G mutation in the viral spike protein, a SNP which is thought to increase the viral load in infected patients and has rapidly become the dominant SARS-CoV-2 strain during the COVID-19 pandemic^27^. We tested our designs in a one-pot SNP-LAMP assay using *in vitro* transcribed RNA fragments for the wild-type and mutant spike protein variants (Figs. 4ab). Our SNP-LAMP assay readily distinguished the D614G spike RNA from wild-type (p=5.8*10^−5^). Though its fluorescence was much lower than the D614G variant, the wild-type spike RNA could also be distinguished from background (p=0.0041). These results indicate that our SNP-LAMP assay can simultaneously detect general SARS-CoV-2 infection, in addition to the presence of a specific viral strain.

**Figure 4.**
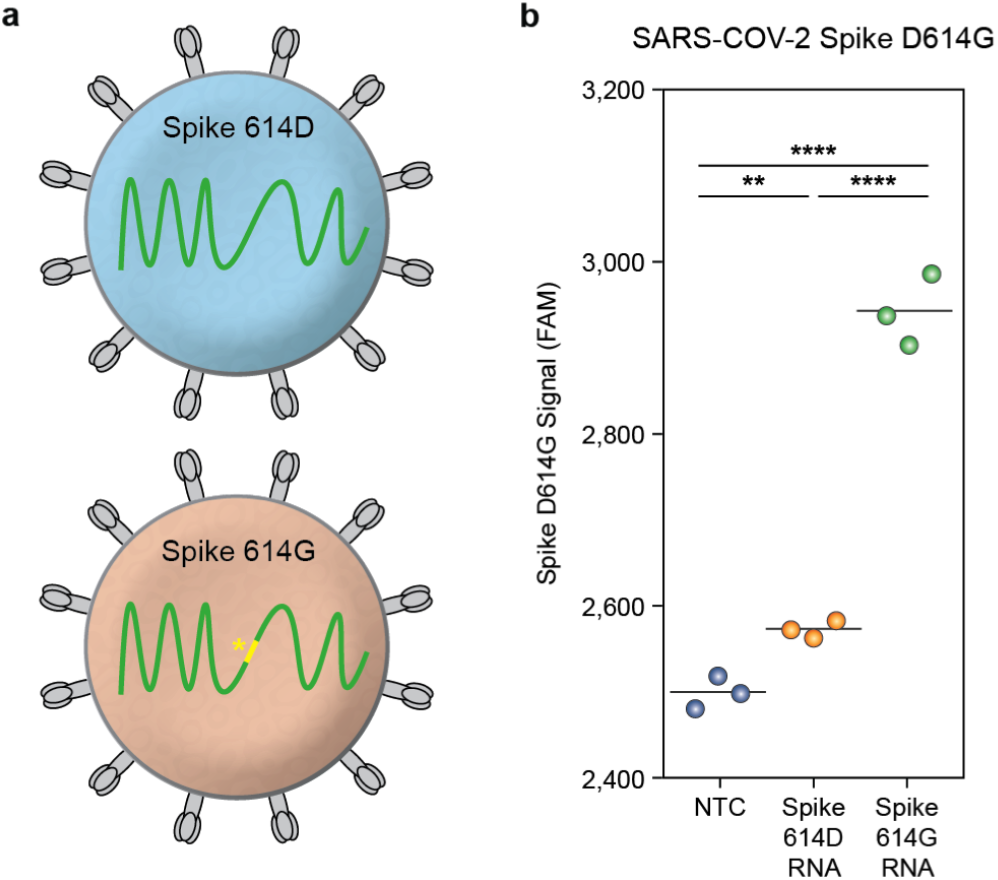
A SNP-LAMP assay for SARS-CoV-2 strain identification. (**a**) Two prevalent SARS-CoV-2 strains in circulation during the COVID-19 global pandemic. The D614G SNP in the spike protein is thought to confer increased viral load. (**b**) Endpoint results for SARS-CoV-2 spike D614G SNP probe after LAMP and annealing. As expected, the probe reliably distinguished a 614G spike RNA fragment from a 614D fragment (p=5.8*10^−5^) and a non-template control (p=3.7*10^−5^).

## Discussion

Single Nucleotide Polymorphisms (SNPs) comprise the majority of genetic variations between individuals^1^ and have implications in human disease^1^, pathogen antibiotic resistance^7^, and agricultural production^6,8^. Detecting SNPs is challenging due to their high similarity with non-SNP sequences, and full DNA sequencing is the most reliable and commonly used method to identify SNPs. DNA sequencing methods require advanced sample preparation and expensive instrumentation, and thus cannot be easily deployed for on-site or low-resource testing^9^. In this work we have developed a simple one-pot method to distinguish complex nucleic acid samples that differ by only a single base. Our approach leverages LAMP-based target amplification and competitive “sink” DNA strands to favor specific activation by SNP products. We demonstrated the robustness of our SNP-LAMP assay by detecting three distinct SNPs in highly complex total RNA samples. We also developed a simple and streamlined assay to detect SARS-CoV-2 strains that could be deployed on a large scale and in low-resource settings.

Previous LAMP-based SNP detection methods are based on either primer mismatches or SNP-specific fluorescent probes. These strategies can be unreliable due to promiscuous polymerase mismatch extension activity^18^, which results in false positives, limited ability to resolve signal differences between highly similar SNP and non-SNP targets, and constraints in designing primer/probe sets that restrict the SNP loci that can be targeted. We were not able to get either of these approaches to work in our hands despite trying a total of 3 different designs. Our competitive SNP-LAMP approach overcomes these limitations by using competing sink strands to drastically enhance the specificity for the target SNP. This resulted in a highly reliable SNP detection method that worked on the first attempt for all targets tested without additional optimization.

We developed a computational pipeline to design competitive SNP-LAMP probes with high sensitivity and specificity. The possible probe space is massive and very few meet our SNP detection criteria. To traverse this design space, we employed a hybrid genetic algorithm (GA) and hill-climbing approach to optimize a probe set’s thermodynamic properties. When initialized from ten random starting states, this algorithm consistently converged to highly specific and sensitive designs. Although we didn’t perform a rigorous head-to-head comparison, we did observe that our computationally designed probe sets had superior SNP:non-SNP specificity relative to manually designed probes. This result is expected since computational optimization can search a much larger design space than human rational design. We believe our computational probe design method can be readily generalized to target any possible SNP of interest.

Our one-pot SNP-LAMP method can rapidly detect SNPs with simple protocol that could be performed by individuals with minimal laboratory training and equipment. The nucleic acid sample is added to a tube containing premixed detection probes, LAMP primers, and a LAMP master mix. This sample is then heated at 65 C for 60-75 minutes and the probes are hybridized by annealing from 95 C to 21 C over 37 minutes. The fluorescence of the sample is measured to provide an assay result in approximately 2 hours. The inexpensive cost of our assay is also a major advantage for large-scale testing. While sanger sequencing assays typically cost $4-7 USD per reaction, the LAMP protocols used here consume less than $1 USD in reagents.

Some applications may require detecting SNPs at low variant allele frequencies (VAF). These include rare dominant active mutations found in genes with high copy numbers or also mixed samples from multiple individuals. We demonstrated that the *TP53* and *NOTCH1* probes could detect their target sequence at SNP frequencies below 1%. However, these experiments were performed DNA oligos and the results may not generalize to actual SNP-LAMP assays. LAMP is a stochastic process with exponential kinetics, and the final LAMP products may not reflect the original variant frequencies within the sample. In cases where SNPs must be detected at low VAFs, a linear amplification method such as rolling circle amplification (RCA)^28^ may provide more reliable frequency estimates.

As we were developing our competitive SNP-LAMP method, we found the competitive probes could cause inhibition of LAMP-based target amplification. We found one-pot SNP-LAMP reactions could only include up to 100 nM of each probe and sink strand before inhibition caused an issue. This constraint limits the total fluorophore in the system and therefore maximum fluorescence output that can be achieved. The fluorescence signal is still easily detectable on a standard laboratory qPCR instrument; however, the signal:background ratio is only ~1.1. We found that increased LAMP primer concentrations could improve the amplification reaction rate and help to overcome inhibition by the probe strands.

Our SNP-LAMP method is simple, rapid, low-cost, and can be performed on basic laboratory equipment. There are several additional modifications that would enable a true point-of-care assay that could be deployed in the field or other low-resource settings. Our current SNP-LAMP assay involves multiple incubation temperatures that require a temperature-adjustable heating device. Our method’s temperature-annealing step is necessary to bring the reaction to thermodynamic equilibrium because the probes were designed based on thermodynamics. Isothermal detection schemes could be devised by considering hybridization kinetics and incorporating single-stranded ‘toehold’ sequences to direct probe binding via DNA branch migration^29^. Our assay also relies on a fluorescent readout that can be challenging to perform on site. There are other label-free detection methods that rely on simple DNA hybridization and strand displacement to generate electrical readouts^30,31^. In theory our SNP-LAMP strategy could be adapted to a have a simple electrical readout.

Field-testing applications, as well as point-of-care assays in developing regions, could also benefit from eliminating cold supply chain requirements. LAMP reagents can be lyophilized and deployed at room temperature^32^, and even packaged in pre-made reaction tubes which only require a liquid sample to be added^33^. Since our method does not rely on pre-annealing the probe and sink complexes before the reaction, they could simply be lyophilized with the other LAMP reagents. We also believe our method may work well on crude nucleic acid samples that have not been processed or purified. Both LAMP and DNA hybridization processes are extremely tolerant to cellular debris, additives, and inhibitors^34,35^. In this case a crude sample could simply be added to lyophilized assay reagents to provide a streamlined workflow that requires minimal hands-on processing and laboratory equipment.

Single nucleotide polymorphisms are crucial drivers of many biological processes, with important implications in human diseases ranging from cystic fibrosis^36^ to cancer^1^. SNPs can also contribute to other undesirable phenomena such as antibiotic resistance in microbes^7^ and agricultural breeding issues^6,8^. Competitive SNP-LAMP provides a reliable, simple, rapid, and low cost SNP detection method that could be deployed for on-site testing in the field or in developing areas of the world^15^. This method will empower researchers across the life sciences by providing a universal solution for point-of-care SNP detection.

## Materials and Methods

### LAMP Primer Design

We first identified SNP targets in target cell lines based on data from the Broad Institute Cancer Cell Line Encyclopedia^37^. We then used the genomic location to retrieve target non-SNP (wild-type) and SNP sequences from the NCBI genome browser^38^. For each gene target, we designed LAMP primers to target these sequence regions using PrimerExplorer V5 software^39^ (Eiken Chemical Co.), placing the SNP base within the LAMP dumbbell loop structure.

### Thermodynamic Simulations of Probe Set Binding

We predicted the equilibrium concentration of each DNA complex using the *complexes* and *concentrations* functions from the NUPACK software package^25^. For each probe set, we simulated three conditions: one with 1μM of a linear DNA strand representing the SNP target sequence, one with 1μM of the non-SNP sequence, and a background condition with no non-SNP or SNP target present. For simplicity, we assumed that 1μM of each probe and sink strand is present in each simulation, as well as 65mM NaCl and 8mM MgCl_2_, matching our SNP-LAMP conditions. We then calculated the percentage of free probeF strand in each simulation to predict the fitness of each probe using the following equation:

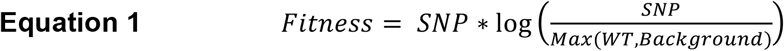

### Probe Design Space Simulations

We simulated 10,000 probe sets by first randomly generating 10,000 pairs of SNP and target sequence pairs. We then created a probe set for each target sequence by truncating a random number of bases from each terminus of the SNP sequence and its complement, and the non-SNP sequence and its complement. We ensured that every complex in each probe set had a duplex at least 6 bases long, and that the probe duplex contained a blunt end for fluorophore and quencher placement. We then performed the thermodynamic simulations described above to predict the fitness of each probe using equation 1, listed above.

### Approximate Tm Calculations

We performed approximate Tm calculations for each DNA duplex in order to screen out highly unfit probes before more computationally intensive thermodynamic simulations. For duplexes greater than 13 bases in length, we used a formula derived by Wallace et al.^40^:

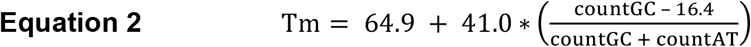

For duplexes less than or equal to 13 bases in length, we used the Marmur rule^41^:

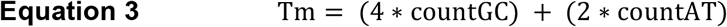

### Principal Component Analysis of Probe Design Space

We performed principal component analysis using the sci-kit-learn package for Python 3.3^42^. We calculated a set of 12 features for each probe representing the binding affinity of the probe and sink complexes using the approximate Tm calculations described above, along with several other relevant design aspects. Features are listed in Table S1. We fit our PCA to the 10,000 random simulated probe sets we generated, and reduced these 12 features to 2 principal components for plotting and for probe screening in our algorithm.

### Computational Probe Design using Genetic Algorithm and Hill Climbing

The genetic algorithm starts by generating an initial population of 128 probe sets that contain a random number of complementary bases on either side of the SNP base position. We include additional constraints that the probe duplex must be greater than 6 bases in length, containing a blunt end to accommodate the fluorophore:quencher pair, and that the bottom strands of both the probe and sink duplexes are completely covered by their complement. This initial population is then evolved by (1) filtering candidate designs to remove designs that do not occupy the high-fitness region of the PCA space (Fig. S1), (2) evaluating each member’s fitness using NUPACK^25^ and our design objective function, (3) selecting the top ranked probes as ‘parents’ for the next generation, (4) randomly crossing parents by selecting a probe complex from one and a sink complex from the other to generate ‘children’, (5) mutating the children by randomly adding/removing a single terminal base to generate new population, and (6) repeating steps 1-5 over 7 generations, halving the population size at each generation. After the genetic algorithm optimization, we identify the top design and perform hill climbing to exhaustively search the local design space and ensure we have reached a maxima. This final probe design should be highly optimized for SNP detection, with a high specificity and signal.

In each hill-climbing iteration, we generated 16 possible mutant probe sets by adding or removing 1 base from each terminus of each probe and sink strand. We performed NUPACK thermodynamic simulations on each mutant probe set and calculated its fitness as in our genetic algorithm. We then moved to the mutant probe set with the greatest fitness gain over the current probe, and repeated the hill-climbing algorithm. When none of the mutant probe sets had a greater fitness than the current one, we returned the current probe set as the final probe set design.

Upon completion of our algorithm, we retrieved the fittest probe in the final generation, and added a fluorophore and quencher to a blunt end of the probe duplex. We then added poly-T tails to all unmodified 3’ termini in the probe and sink complexes in order to prevent polymerase extension on the LAMP product.

### DNA Complexes and Primer Mixes

We ordered all DNA oligos from Integrated DNA Technologies (Coralville, Iowa), and dissolved each into nuclease-free water (Thermo Fisher) prior to storage at -20 °C. We prepared LAMP primer mix and probe set mix stocks in nuclease-free water (Thermo Fisher) and stored at -20 °C. On the day of experiment, we thawed each mixture on ice while protecting from light, and subsequently added to LAMP reactions.

### Production of mRNAs using in vitro transcription

DNA templates for SARS-CoV-2 614D and 614G fragments were synthesized by Integrated DNA Technologies (Coralville, Iowa), each containing an upstream T7 promoter. We PCR amplified each fragment using Phusion® High-Fidelity DNA Polymerase (New England Biolabs) and purified the resulting amplicons with the DNA Clean & Concentrator-5 kit (Zymo Research). We then performed *in vitro* transcription from these amplicons using the HiScribe™ T7 High Yield RNA Synthesis Kit (New England Biolabs), and purified the resulting RNA using a GeneJET RNA Purification Kit (Thermo Scientific). We quantified each RNA sample’s concentration using a NanoDrop™ Spectrophotometer (Thermo Scientific) and stored RNA stocks at -80 °C in RNase-free water. Primer and DNA fragment sequences are given in Table S2.

### Total RNA Extraction from Cell Lines

We subcultured MOLT-4 cells (American Type Culture Collection) in a 1:8 ratio every two days in RPMI-1640 Medium (Gibco) supplemented with 10% FBS (Gibco) and 1X Antibiotic-Antimycotic (Gibco). We subcultured SK-BR3 cells (American Type Culture Collection) in a 1:4 ratio every two days in DMEM, high glucose (Gibco) supplemented with 10% FBS and 1X Antibiotic-Antimycotic. We collected approximately 2.5 million cells of each type and purified their total RNA using a GeneJET RNA Purification Kit (Thermo Scientific). We quantified the concentration of each RNA sample using a NanoDrop™ Spectrophotometer (Thermo Scientific) and stored RNA stocks at -80 °C in RNase-free water.

### SNP-LAMP and Annealing Assays

We performed SNP-LAMP and annealing assays in triplicate using a Bio Rad CFX Connect qPCR machine. Except in anneal-only experiments, we incubated the reactions at 65 °C to allow LAMP amplification while monitoring FAM, HEX, and/or SYBR fluorescence channels. We subsequently heated the reaction to 95°C for two minutes. We then annealed the probes by lowering the temperature by 1°C every 30 seconds and monitored FAM or HEX fluorescence channels at each step. Each reaction comprised a total volume of 10μL, consisting of 1X WarmStart LAMP Master Mix (New England Biolabs), and 0.5 U/μL SUPERase•In™ RNase Inhibitor (Invitrogen), with primer concentrations given in Table S3 and probe concentrations given in Table S4. Primer, target oligo, and probe set sequences are given in Table S2. LAMP durations and input RNA amounts are listed in Table S3, while probe and target oligo concentrations are listed in Table S4. In cases where no target oligo or RNA was present, we added water as a non-template control.

### Sanger Sequencing of LAMP Amplicons

Using the *MT-CO2, ACTB, TP53*, and *NOTCH1* LAMP primers listed in Table S2, we performed LAMP amplification from 10ng of MOLT-4 or SK-BR-3 total RNA in 25μL reactions for 90 minutes, as described above. We verified reaction completion by eye using turbidity^43^. Upon reaction completion, we added 10μg of RNase A (Thermo Fisher) and incubated at 37°C for 15 minutes to destroy cellular RNA. We then purified products from each LAMP reaction using a Zymo DNA Clean & Concentrator-5 kit, eluting in nuclease-free water. We quantified each product using a NanoDrop™ Spectrophotometer (Thermo Scientific) and submitted for sanger sequencing using the product’s corresponding FIP and BIP primers listed in Table S2. In the case of *MT-CO2*, the primers used for sequencing differed from those used in SNP-LAMP experiments, as we later found a LAMP primer set with superior reaction speed. However, both primer sets targeted the same SNP mutation.

### Statistical Testing

To obtain the p-values reported in this work, we first performed a two-tailed t-test to verify that a significant effect is present. When the p-value for this test was above 0.05, we reported it directly. Otherwise, we performed a one-sided t-test and reported the p-value. We performed all tests with n=3, assuming homoscedasticity.

## Data and Code Availability

Data and code that support the findings of this study are available from the corresponding author upon reasonable request.

## Acknowledgements

P.A.R. is a Damon Runyon-Rachleff Innovator supported (in part) by the Damon Runyon Cancer Research Foundation (DRR-40-16).

## Author Contributions

L.B.H. and P.A.R. conceived the project. L.B.H. and C.R.C. performed the experiments. L.B.H. analyzed the data with feedback from P.A.R.. L.B.H. and P.A.R. wrote the manuscript.

## Competing Interests

The authors declare no competing interests.

## Notes

### Competing Interest Statement

The authors have declared no competing interest.

