## Supplemental information for "Competitive SNP-LAMP probes for rapid and robust single-nucleotide polymorphism detection"

### Supplementary Information

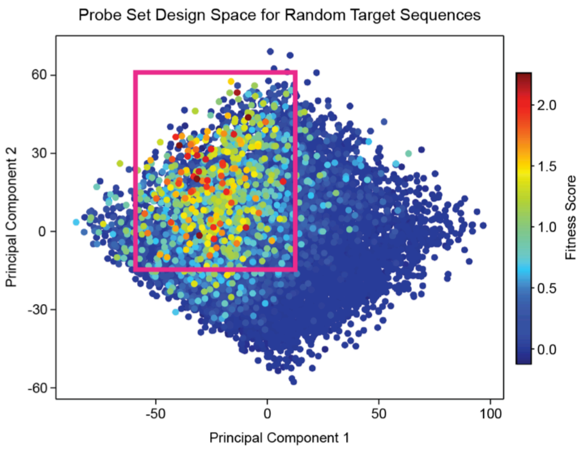


**Figure S1 |** SNP probe design space visualized after PCA. To improve the convergence of our optimization algorithm, we initialized all newly generated probes within the highest fitness space (magenta).

| | | Target Sequences | Sequencing Reads | |
|---|---|---|---|---|
| *MT-CO2* c.186A>G | **Wildtype** | AAATAGA A ACCGTCT | AAATAGA A ACCGTCT | SK-BR-3 |
| | **SNP** | AAATAGA G ACCGTCT | AAATAGA G ACCGTCT | MOLT-4 |
| *TP53* c.524G>A | **Wildtype** | GTGAGGC G CTGCCCC | GTGAGGN G CTGCCCC | MOLT-4 |
| | **SNP** | GTGAGGC A CTGCCCC | GTGAGGN A CTGCCCC | SK-BR-3 |
| *NOTCH1* c.4799T>C | **Wildtype** | CGCGTGC T GCACACC | CGCGTGC T GCACACN | SK-BR-3 |
| | **SNP** | CGCGTGC C GCACACC | CGCGTNC C GCACACC | MOLT-4 |

**Figure S2 |** Expected and observed sanger sequencing reads from *MT-CO2*, *TP53*, and *NOTCH1* LAMP amplicons. We used either SK-BR-3 or MOLT-4 cell total RNA as input. In all cases, amplicons contained the expected SNP or non-SNP (WT) base in the loop region of their dumbbell structure.

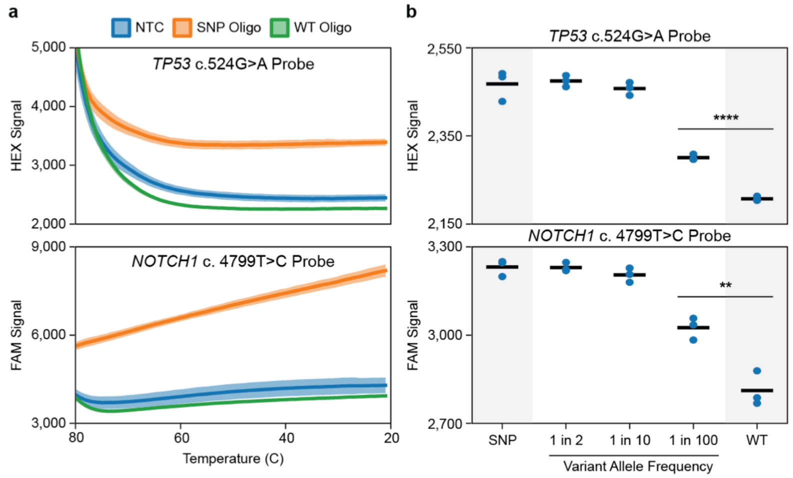


**Figure S3 |** TP53 C.524G>A and NOTCH1 C.4799T>C SNP probes are highly selective. (**a**) Annealing curves for the *TP53* and *NOTCH1* SNP probes. Each was highly selective for a DNA oligo carrying its SNP target sequence over a non-SNP (WT) oligo. (**b**) *TP53 C.524G>A* and *NOTCH1* C.4799T>C probe activation in response to low variant allelic frequencies of the SNP target. Each probe displayed a SNP/non-SNP selectivity of at least 100-fold ($p= 2.3*10^{-5}$ and $p=3.1*10^{-3}$ for *TP53* and *NOTCH1*, respectively).

**Table S1.** Probe set features used in principal component analysis, and their coefficients in each component.

| **Complex Tms** | **Component 1** | **Component 2** |
|---|---|---|
| ProbeF:ProbeQ Tm | -0.2457 | 0.3782 |
| ProbeF:SNP Tm | -0.3275 | 0.5221 |
| Sink:Sink* Tm | 0.2548 | 0.4251 |
| Sink:WT Tm | 0.2948 | 0.5536 |

| **Tm Differences** | | |
|---|---|---|
| ProbeF:ProbeQ Tm - Sink:Sink* Tm | -0.5005 | -0.0469 |
| ProbeF:SNP Tm - Sink:WT Tm | -0.6223 | -0.0316 |
| ProbeF:SNP Tm - ProbeF:ProbeQ Tm | -0.0818 | 0.1439 |
| Sink:WT Tm - Sink:Sink* Tm | 0.0400 | 0.1285 |

| **SNP Base Proximity to Termini** | | |
|---|---|---|
| ProbeF SNP Distance | -0.1091 | 0.1470 |
| ProbeQ SNP Distance | -0.0731 | 0.0823 |
| Sink SNP Distance | -0.1091 | 0.1470 |
| Sink* SNP Distance | -0.0731 | 0.0823 |

**Table S2.** DNA sequences of LAMP primers, probe sets, gBlocks, and DNA target oligo controls.

| ***ACTB* C.776A>C SNP-LAMP Probe Set** | |
|---|---|
| *ACTB* C.776A>C ProbeF | **6-FAM**-CGAAGGCTGGAAGAGTGCCGCAGGGCAGCGTTTTTTTT |
| *ACTB* C.776A>C ProbeQ | TGCGGCACTCTTCCAGCCTTCG-**IowaBlackFQ** |
| *ACTB* C.776A>C Sink | AAGGCTGGAAGAGTGCCTCAGGGCAGCG |
| *ACTB* C.776A>C Sink* | TGAGGCACTCTTCCAGCCTT |
| ***ACTB* C.776A>C Target Sequence Oligos** | |
| *ACTB* C.776A>C WT Target Oligo | CGCTGCCCTGAGGCACTCTTCCAGCCTT |
| *ACTB* C.776A>C SNP Target Oligo | CGCTGCCCTGCGGCACTCTTCCAGCCTT |

| ***MT-CO2* C.186A>G LAMP Primers (SNP-LAMP)** | |
|---|---|
| *MT-CO2* C.186A>G F3 - 1 | TTTTCCTAACACTCACAACAA |
| *MT-CO2* C.186A>G B3 - 1 | TCGTAGGTTCAGTACCATTG |
| *MT-CO2* C.186A>G FIP - 1 | GAGGACTAGGATGATGGCGGCTAACATCTCAGACGCTCA |
| *MT-CO2* C.186A>G BIP - 1 | CTCCCATCCCTACGCATCCTGCCAATTGATTTGATGGTAAGG |
| *MT-CO2* C.186A>G LB - 1 | TTACATAACAGACGAGGTCAACGAT |
| ***MT-CO2* C.186A>G LAMP Primers (Sanger sequencing)** | |
| *MT-CO2* C.186A>G F3 - 2 | ACTTCCCCTATCATAGAAGAG |
| *MT-CO2* C.186A>G B3 - 2 | CTCGTCTGTTATGTAAAGGATG |
| *MT-CO2* C.186A>G FIP - 2 | GGCATACAGGACTAGGAAGCAGTCACCTTTCATGATCACGC |
| *MT-CO2* C.186A>G BIP - 2 | AATACTAACATCTCAGACGCTCAGGGCGATGAGGACTAGGATGA |
| ***MT-CO2* C.186A>G SNP-LAMP Probe Set** | |
| *MT-CO2* C.186A>G ProbeF | **HEX**-TGGCGGGCAGGATAGTTCAGACGGTCTCTATTTTTTTTT |
| *MT-CO2* C.186A>G ProbeQ | AGACCGTCTGAACTATCCTGCCCGCCA-**IowaBlackFQ** |
| *MT-CO2* C.186A>G Sink | TGGCGGGCAGGATAGTTCAGACGGTTTCTATTTTTTTT |
| *MT-CO2* C.186A>G Sink* | AAATAGAAACCGTCTGAACTATCCTGTTTTTT |
| ***MT-CO2* C.186A>G Target Sequence Oligos** | |
| *MT-CO2* C.186A>G WT Target Oligo | AAATAGAAACCGTCTGAACTATCCTGCCCGCCA |

| | |
|---|---|
| *MT-CO2* C.186A>G SNP Target Oligo | AAATAGAGACCGTCTGAACTATCCTGCCCGCCA |

| | |
|---|---|
| ***TP53 C.524G>A* LAMP Primers** | |
| *TP53 C.524G>A* F3 | TTGCCAACTGGCCAAGAC |
| *TP53 C.524G>A* B3 | CGCAAATTTCCTTCCACTCG |
| *TP53 C.524G>A* FIP | ACTGCTTGTAGATGGCCATGGCCTGTGCAGCTGTGGGTTG |
| *TP53 C.524G>A* BIP | CAGCACATGACGGAGGTTGTGAGGGCCAGACCATCGCTAT |
| *TP53 C.524G>A* LF | ACGCGGGTGCCGG |
| ***TP53 C.524G>A* SNP-LAMP Probe Set** | |
| *TP53 C.524G>A* ProbeF | **HEX**-GGCACTGCCCCCACCATGAGCGCTGCTCTTTTTTTT |
| *TP53 C.524G>A* ProbeQ | AGCAGCGCTCATGGTGGGGGCAGTGCC-**IowaBlackFQ** |
| *TP53 C.524G>A* Sink | GGCGCTGCCCCCACCATGAGCGCTGCTTTTTTTT |
| *TP53 C.524G>A* Sink* | TCATGGTGGGGGCAGCGTTTTTTTT |
| ***TP53 C.524G>A* Target Sequence Oligos** | |
| *TP53 C.524G>A* WT Target Oligo | CTGAGCAGCGCTCATGGTGGGGGCAGCGCC |
| *TP53 C.524G>A* SNP Target Oligo | CTGAGCAGCGCTCATGGTGGGGGCAGTGCC |

| | |
|---|---|
| ***NOTCH1* C.4799T>C LAMP Primers** | |
| *NOTCH1* C.4799T>C F3 | GCAACAGCGCGGAGTG |
| *NOTCH1* C.4799T>C B3 | GGAAGATCATCTGCTGGCC |
| *NOTCH1* C.4799T>C FIP | GCATCAGCACCACCACCACCGTGCGGAGCATGTACCCG |
| *NOTCH1* C.4799T>C BIP | CGCAACAGCTCCTTCCACTTCCTGCGTCACGCTTGAAGAC |
| *NOTCH1* C.4799T>C LF | CCGGCCGCCAGCCT |
| ***NOTCH1* C.4799T>C SNP-LAMP Probe Set** | |
| *NOTCH1* C.4799T>C ProbeF | **6-FAM-** GCGGGAGCTCAGCCGCGTGCCGCACACCAACGTGTTTTTTTT |
| *NOTCH1* C.4799T>C ProbeQ | GTTGGTGTGCGGCACGCGGCTGAGCTCCCGC-**IowaBlackFQ** |
| *NOTCH1* C.4799T>C Sink | TGCGGGAGCTCAGCCGCGTGCTGCACACCAACGTTTTTTTT |
| *NOTCH1* C.4799T>C Sink* | GTGCAGCACGCGGCTGAGCTTTTTTTTT |
| ***NOTCH1* C.4799T>C Target Sequence Oligos** | |
| *NOTCH1* C.4799T>C WT Target Oligo | CACGTTGGTGTGCAGCACGCGGCTGAGCTCCCGCA |
| *NOTCH1* C.4799T>C SNP Target Oligo | CACGTTGGTGTGCGGCACGCGGCTGAGCTCCCGCA |

| | |
|---|---|
| **SARS-CoV-2 Spike D614G LAMP Primers** | |
| Spike D614G F3 | TGCTGACACTACTGATGC |
| Spike D614G B3 | GTAGAATAAACACGCCAAGT |
| Spike D614G FIP | ACTGACACCACCAAAAGAACATGTGTCCGTGATCCACAGAC |
| Spike D614G BIP | AATACTTCTAACCAGGTTGCTGTTCGAGTAAGTTGATCTGCATGAA |
| Spike D614G LF | AATGTCAAGAATCTCAAGT |
| **SARS-CoV-2 Spike D614G SNP-LAMP Probe Set** | |
| Spike D614G ProbeF | ATCAGGGTGTTAACTGCACAGAAGTCCCTG**-6-FAM** |
| Spike D614G ProbeQ | **IowaBlackFQ-** CAGGGACTTCTGTGCAGTTAACACCCTGAAAAAAAA |
| Spike D614G Sink | TCAGGATGTTAACTGCACAGAAGTCCCAAAAAAAA |
| Spike D614G Sink* | TTCTGTGCAGAAAAAAAA |
| **SARS-CoV-2 Spike D614G PCR Primers** | |
| T7 Promoter Forward | TAATACGACTCACTATAGGG |
| Spike PCR Reverse | ACATTAGAACCTGTAGAATAAAC |
| **SARS-CoV-2 Spike D614G gBlock Fragments** | |
| SARS-CoV-2 Spike 614D Fragment gBlock | TAATACGACTCACTATAGGGGAATTGTGAGCGGATAACAATTCCCCTCTAGAAATAATTTTGTTTAACTTTAAGAAGGAGATATACATATGGGCAGAGACATTGCTGACACTACTGATGCTGTCCGTGATCCACAGACACTTGAGATTCTTGACATTACACCATGTTCTTTTGGTGGTGTCAGTGTTATAACACCAGGAACAAATACTTCTAACCAGGTTGCTGTTCTTTATCAGGATGTTAACTGCACAGAAGTCCCTGTTGCTATTCATGCAGATCAACTTACTCCTACTTGGCGTGTTTATTCTACAGGTTCTAATGT |
| SARS-CoV-2 Spike 614G Fragment gBlock | TAATACGACTCACTATAGGGGAATTGTGAGCGGATAACAATTCCCCTCTAGAAATAATTTTGTTTAACTTTAAGAAGGAGATATACATATGGGCAGAGACATTGCTGACACTACTGATGCTGTCCGTGATCCACAGACACTTGAGATTCTTGACATTACACCATGTTCTTTTGGTGGTGTCAGTGTTATAACACCAGGAACAAATACTTCTAACCAGGTTGCTGTTCTTTATCAGGGTGTTAACTGCACAGAAGTCCCTGTTGCTATTCATGCAGATCAACTTACTCCTACTTGGCGTGTTTATTCTACAGGTTCTAATGT |

**Table S3.** Primer concentrations, RNA input concentrations, and LAMP duration for each experiment.

| Experiment | Format | [Primers]* | [RNA] | LAMP Duration |
|---|---|---|---|---|
| *ACTB* Probe | Anneal only | Standard | - | - |
| *MT-CO2* Probe | One-pot | Standard | 1ng/μL | 75 minutes |
| *NOTCH1* Probe | One-pot | Double | 2ng/μL | 75 minutes |
| *TP53* Probe | One-pot | Double | 1ng/μL | 75 minutes |
| *NOTCH1* V.A.F. | Anneal only | - | - | - |
| *TP53* V.A.F. | Anneal only | - | - | - |
| SARS-CoV-2 Spike D614G Probe | One-pot | Standard | 1nM | 60 minutes |

*Standard LAMP primer concentrations consist of 1.6 μM of each FIP/BIP primer, 0.2 μM of each F3/B3 Primer, and 0.4 μM of each LoopF/LoopB Primer

**Table S4.** Probe, sink, and target oligo concentrations used in each experiment.

| Experiment | Format | [ProbeF] | [ProbeQ] | [Sink] | [Sink*] | [WT oligo] | [SNP oligo] |
|---|---|---|---|---|---|---|---|
| *ACTB* Probe | Anneal only | 1μM | 1μM | 1μM | 1μM | 1μM | 1μM |
| *MT-CO2* Probe | One-pot | 100nM | 100nM | 100nM | 100nM | - | - |
| *NOTCH1* Probe | One-pot | 100nM | 100nM | 100nM | 100nM | - | - |
| *TP53* Probe | One-pot | 100nM | 100nM | 100nM | 100nM | - | - |
| *NOTCH1* V.A.F. | Anneal only | 100nM | 100nM | 10μM | 10μM | 0-10μM | 0-100nM |
| *TP53* V.A.F. | Anneal only | 100nM | 100nM | 10μM | 10μM | 0-10μM | 0-100nM |
| SARS-CoV-2 Spike D614G Probe | One-pot | 100nM | 100nM | 100nM | 100nM | - | - |
